# The ColR/S two-component system is a conserved determinant of host association across *Pseudomonas* species

**DOI:** 10.1101/2021.12.14.472530

**Authors:** Christina L. Wiesmann, Yue Zhang, Morgan Alford, David Thoms, Melanie Dostert, Andrew Wilson, Daniel Pletzer, Robert E. W. Hancock, Cara H. Haney

**Author notes:** These authors contributed equally to this work (Christina L. Wiesmann, Yue Zhang). Author order was determined alphabetically.

## Abstract

Members of the bacterial genus *Pseudomonas* form mutualistic, commensal and pathogenic associations with diverse hosts. The prevalence of host association across the genus suggests that symbiosis may be a conserved ancestral trait and that distinct symbiotic lifestyles may be more recently evolved. Here we show that the ColR/S two-component system, part of the *Pseudomonas* core genome, is functionally conserved between *Pseudomonas aeruginosa* and *Pseudomonas fluorescens*. Using plant rhizosphere colonization and virulence in a murine abscess model, we show that *colR* is required for commensalism with plants and virulence in animals. Comparative transcriptomics revealed that the ColR regulon has diverged between *P. aeruginosa* and *P. fluorescens* and deleting components of the ColR regulon revealed strain-specific, but not hostspecific, requirements for ColR-dependent genes. Collectively, our results suggest that ColR/S allows *Pseudomonas* to sense and respond to a host, but that the ColR-regulon has diverged between *Pseudomonas* strains with distinct lifestyles.

## Introduction

Plants and animals are colonized by symbiotic bacteria. These include mutualists that help with nutrient uptake or pathogen protection, commensals that live in association with hosts without causing harm or benefit, and pathogens that cause disease. Interestingly, transitions between symbiotic lifestyles^1,2^ and even hosts^3^ can occur over relatively short evolutionary distances. While mechanisms underlying bacterial virulence and mutualism are widely studied, relatively little research has focused on the mechanisms by which bacteria initiate symbiosis and if these mechanisms are conserved across closely-related pathogenic and commensal bacteria.

Both plants and animals have functionally similar receptors and signaling pathways that allow them to sense and respond to the presence of bacteria^4^. Accordingly, *Pseudomonas aeruginosa* makes use of the same virulence factors required to infect both plants and animals^5,6^ suggesting *P. aeruginosa* targets similar cellular processes across hosts. Consistently, the nitrogen-fixing plant mutualist *Sinorhizobium meliloti* has genes required for symbiosis that are homologous to virulence factors in the animal pathogen *Brucella abortus* and the plant pathogen *Agrobacterium tumefaciens*^3^. Regardless of symbiotic lifestyle, host-associated bacteria must be able to colonize their host. As a result, we hypothesized that genes required for host colonization may be shared by pathogens, commensals, and mutualists.

The bacterial genus *Pseudomonas* includes plant commensals and opportunistic animal and plant pathogens^7–9^ and thus provides a model genus to study conserved symbiotic mechanisms. We hypothesize that the ability to be symbiotic is ancestral within the genus *Pseudomonas*, and thus should be encoded by components of the core genome, with further evolution of species or strains leading to distinct lifestyles. Indeed, the *Pseudomonas* accessory genome has previously been shown to encode virulence factors^1,10^ while colonization factors may include components of the core genome^11^.

Two-component systems are common signal-transduction cascades in bacteria that sense environmental cues, including the presence of a host, and cause altered gene expression. Orthologous two-component systems in *Brucella abortus* (BvrR/S), *Sinorhizobium meliloti* (ExoS/ChvI) and *Agrobacterium tumefaciens* (VirA/VirG), are required for virulence in animals and mutualism or virulence in plants^12–15^. The ColR/S two-component system is required for colonization of the tomato rhizosphere by *Pseudomonas fluorescens* WCS365^16,17^, and ColS is required for *Pseudomonas aeruginosa* TBCF10839 virulence during infection of the nematode *Caenorhabditis elegans*^18^. As different species spanning the *Pseudomonas* genus can be host associated^19^, we hypothesized that distantlyrelated *Pseudomonas* species may use conserved two-component signaling pathways to sense and adapt to diverse host environments.

Because ColR/S has been identified in phylogenetically diverse *Pseudomonas* spp. as required for rhizosphere colonization^16,20^ or virulence in *C. elegans*^18^, we tested whether ColR/S, and its regulon, are broadly conserved host-association factors across diverse *Pseudomonas* spp. and diverse hosts. The genes encoding for ColR/S are components of the core *Pseudomonas* genome, and previously identified components of the ColR regulon are broadly conserved across the genus^11^. Collectively this indicates that identifying the genes and mechanisms by which ColR regulates host association may reveal conserved mechanisms that predispose members of the genus *Pseudomonas* for symbiosis.

This study uses *P. fluorescens* WCS365, *P. aeruginosa* PAO1, and *P. aeruginosa* LESB58 as model strains to study the mechanisms of host association across the genus *Pseudomonas. P. fluorescens* WCS365 is a beneficial, plant growth-promoting strain^21^. *P. aeruginosa* PAO1 is a human and plant opportunistic pathogen^22^. It can efficiently colonize the plant rhizosphere, but causes disease in plants including Arabidopsis, sweet basil, canola, and poplar trees^23–25^. *P. aeruginosa* PAO1 also disseminates into the tissue of mice to cause infection, making it an unideal strain to use to study prolonged association with a murine host^26^. *P. aeruginosa* LESB58 was isolated from a cystic fibrosis patient and causes chronic infection in a subcutaneous murine infection model, making it an ideal strain for studying host association over many days^26^. We used growth of *P. aeruginosa* PAO1 and *P. fluorescens* WCS365 in an Arabidopsis rhizosphere model^27^ to identify conserved or divergent *colR-dependent* genes required for rhizosphere colonization. We then used *P. aeruginosa* LESB58 in a murine subcutaneous abscess model to test if the genes required for rhizosphere colonization are also required for virulence in a murine model^26^.

## Results

### The ColR response regulator is highly conserved and required for rhizosphere colonization across *Pseudomonas* species

To assess the degree of functional conservation of the ColR response regulator across the genus, we used the model strains *P. fluorescens* WCS365, *P. aeruginosa* PAO1 and *P. aeruginosa* LESB58 to study parallels in host association. An alignment of the ColR and ColS sequences from six *Pseudomonas* strains from four different species shows a high degree of similarity across the genus (Supplementary Fig. 1).*P. aeruginosa* PAO1 and *P. fluorescens* WCS365 ColR and ColS are 88% and 61% identical at the amino acid level, respectively, suggesting that ColR and ColS are highly conserved between these two species (Supplementary Fig. 1). To test the conservation of the requirement of *colR* in rhizosphere colonization, we deleted c*olR* in *P. fluorescens* WCS365 and in *P. aeruginosa* PAO1. We found that consistent with disruption of *colS* being required for potato root tip colonization^16^, deletion of *colR* leads to a 7-fold decrease in growth of WCS365 (Fig. 1a) and a 10-fold decrease in rhizosphere growth of PAO1 (Fig. 1b) when inoculated into rhizosphere of 12-day old Arabidopsis seedlings grown hydroponically (Methods). Microscopy with *P. aeruginosa* PAO1 expressing GFP revealed lower abundance of the Δ*colR* mutant in the rhizosphere (Fig. 1c). Complementation of *colR* deletion mutants with the copies of the *colR* from their respective strains expressed under the *colR* native promoter restored rhizosphere colonization of the *colR* deletion mutants to wild-type levels or higher (Fig. 1a, b). Because of the high degree of conservation of ColR across species, we tested if *P. aeruginosa* PAO1 *colR* could complement a *P. fluorescens* WCS365 Δ*colR* mutant. We found that PAO1 *colR* expression from a plasmid is able to complement the WCS365 Δ*colR* deletion and rescue rhizosphere growth (Fig. 1a) indicating that *colR* is a conserved rhizophore colonization factor across diverse *Pseudomonas*.

**Figure 1.**
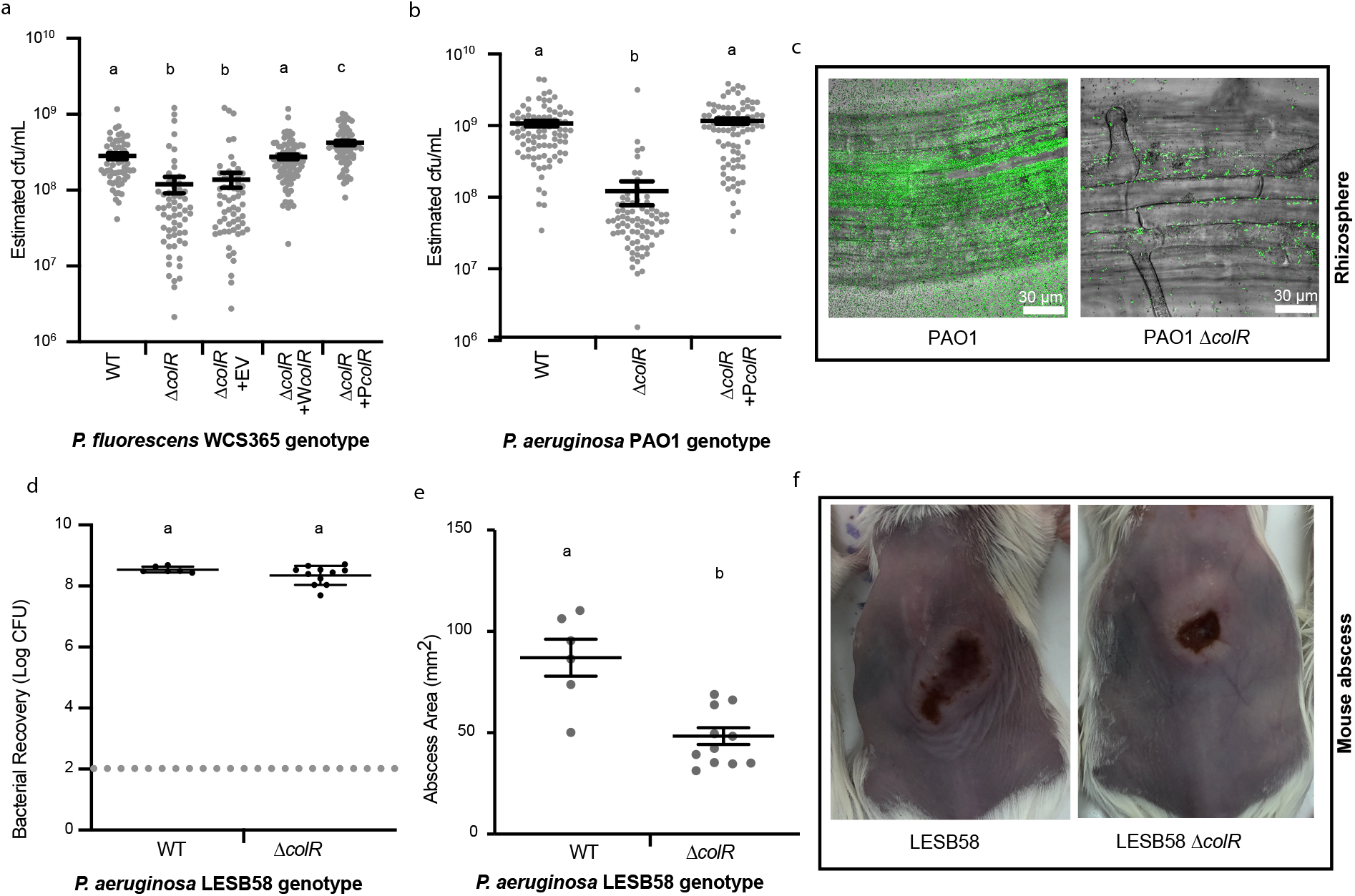
ColR is a conserved host association factor across *Pseudomonas* spp. in plant and murine hosts. **(a)** To test if *colR* is necessary for rhizosphere colonization, seedlings growing hydroponically in 48-well plates were inoculated with GFP-expressing *P. fluorescens* WCS365, WCS365 Δ*colR*, WCS365 Δ*colR* containing an empty vector (EV), a vector expressing WCS365 *colR* under its native promoter (*WcolR*), a vector expressing PAO1 *colR* under its native promoter (*PcolR*) or **(b)** *P. aeruginosa* PAO1, PAO1 Δ*colR*, or PAO1 Δ*colR* expressing *colR* under its native promoter (*PcolR*). **(a and b)** GFP fluorescence values in each plant-containing well were measured 5 days post-inoculation and converted to OD_600_ using a standard curve. Each data point represents the estimated CFU values from a single well containing a single plant. Experiments were repeated three independent times with 10-30 plants per replicate (n = 30-90). Mean +/− standard error is shown and letters indicate significant (P < 0.05) differences as determined by a one-way ANOVA follow by a post-hoc Tukey HSD test. **(c)** Light microscopy of *P. aeruginosa* PAO1 expressing GFP (green) growing for two days on Arabidopsis roots shows that deletion of *colR* leads to visibly less PAO1 colonization of the root. **(d and e)** Wild-type *P. aeruginosa* LESB58 and a LESB58 Δ*colR* mutant were injected (~5 x 10^7^ CFU inoculum) into the subcutaneous thin skeletal muscle on the dorsum of mice. CFU counts **(d)** and abscess lesion size **(e)** were determined 3 days post infection. Means +/− standard error are shown; each dot represents the results from one animal. Different letters indicate significant differences at P < 0.05 determined using a one-way ANOVA followed by a Tukey’s HSD test, or using a t-test if only one comparison was made. **(f)** Representative images of abscess formation by wild-type LESB58 and the Δ*colR* mutant.

Because previous research has shown that virulence mechanisms are conserved across *Pseudomonas aeruginosa* infecting plants and animals^6,28^, we also tested if *colR* was required for virulence in a murine subcutaneous abscess model^26^. We deleted *colR* in the Liverpool Epidemic Strain *P. aeruginosa* LESB58 (Methods; Dataset S1) and found that the *colR* mutant formed a significantly smaller abscess than wild-type LESB58 (54% reduction in size), indicating that *colR* was required for virulence (Fig. 1e, f) In contrast to Δ*colR* mutants in the rhizosphere, the LESB58 Δ*colR* mutant grew to similar levels as wild-type bacteria in an abscess (Fig. 1d) suggesting that *colR* in LESB58 may primarily contribute to virulence rather than growth in the abscess model. Collectively these data indicate that ColR plays a conserved role in host association in both *P. aeruginosa* and *P. fluorescens*.

### The ColR regulons are highly divergent across *Pseudomonas* strains

The requirement of *colR* in symbiosis (commensalism or virulence) in *P. fluorescens* and *P. aeruginosa* led us to hypothesize that *colR* regulates an overlapping set of genes in these two species. To identify ColR-dependent genes in both *P. fluorescens* WCS365 and in *P. aeruginosa* PAO1, we performed RNA-Seq on wild-type and Δ*colR* mutants in the rhizosphere. We grew both PAO1 and WCS365 in the Arabidopsis rhizosphere and in defined media and isolated RNA 6 h post-inoculation to identify rhizosphere-specific *colR*-dependent and independent responses. We chose to use defined M9 minimal media as a control condition to identify genes that are induced in the rhizosphere. RNA from three biological replicates was extracted and sequenced using paired-end 150 bp reads on an Illumina HiSeq. The sequencing reads were mapped to the PAO1 or WCS365 transcriptomes using Salmon^29^. As previously only a draft genome of WCS365 was available, a complete WCS365 genome assembly was generated for use in this study (Methods).

To determine how divergent the transcriptomes were between wild-type bacteria and the *colR* mutants, and between the rhizosphere and M9 minimal media, we performed principal component analysis (PCA). We found that the majority of the differences in gene expression within a single strain (94-97%) were due to growth condition (media vs. rhizosphere) and only a small (1-2%) amount of variance in gene expression was due to bacterial genotype (wild-type vs. Δ*colR*) (Fig. 2a). This suggests that *Pseudomonas* undergo major transcriptional reprogramming in the rhizosphere, but that only a subset of these changes are ColR-dependent.

**Figure 2.**
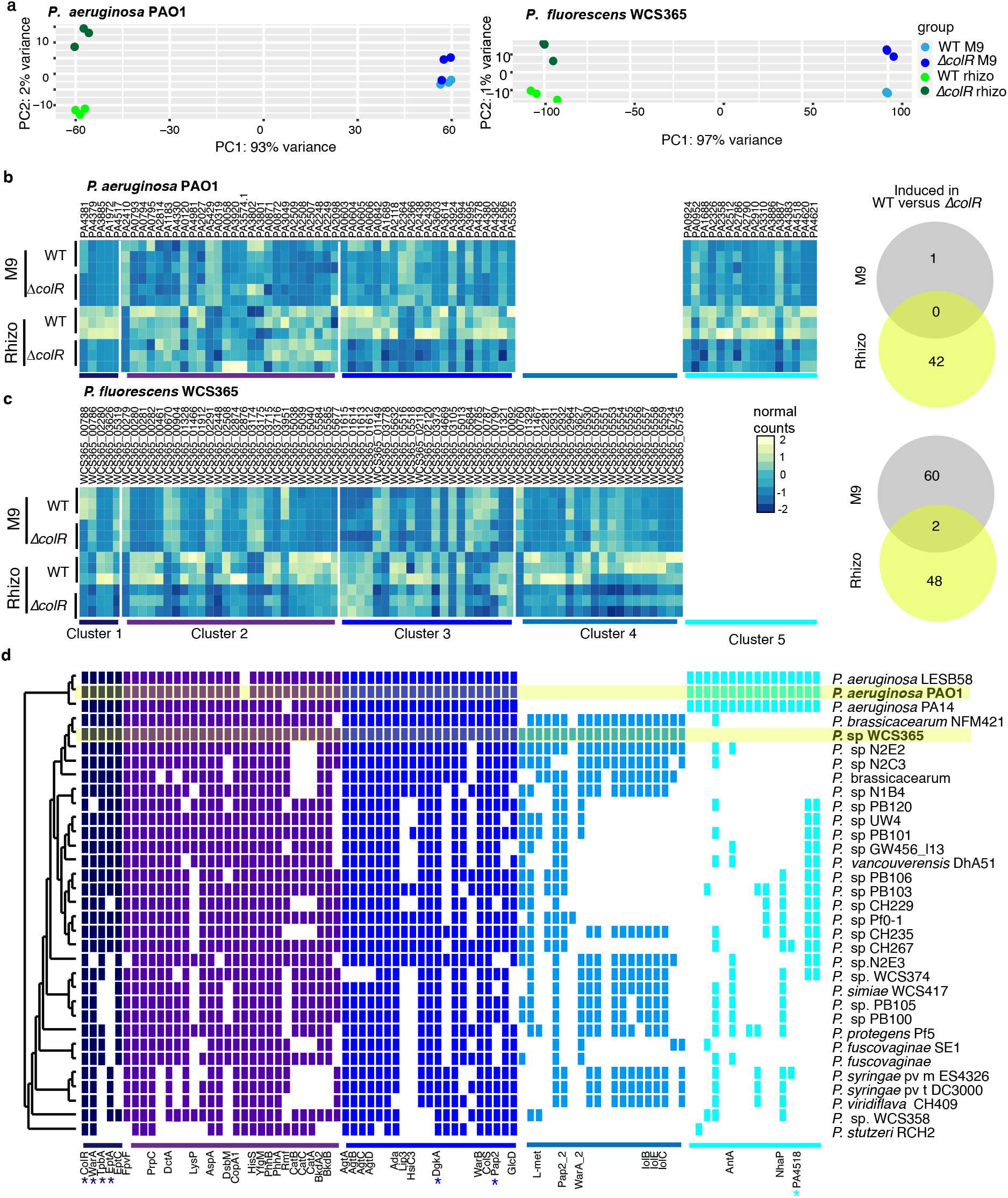
ColR regulates a limited number of genes in the rhizosphere that are distinct between *Pseudomonas* strains. **(a)** PCA plots of transcript count matrices of RNAseq analysis comparing gene expression in wildtype *P. aeruginosa* PAO1 and the PAO1 Δ*colR* mutant (left) or *P. fluorescens* WCS365 and the WCS365 Δ*colR* mutant (right). (**b and c)** Heat maps and Venn diagrams showing the number of significantly differentially expressed genes (greater than 0.585-fold change, padj <0.01) between WT and the *colR* mutant in the rhizosphere or in minimal media in PAO1 **(b)** and WCS365 **(c)**. Gene counts higher than row average are shown in yellow, and gene counts lower than row average are shown in blue. **(d)** A phylogenetic tree showing orthologous genes across *Pseudomonas* strains. Genes fell into 5 distinct clusters: **Cluster 1.** Genes with ColR-dependent expression in both WCS365 and PAO1; **Cluster 2**. Genes with ColR-dependent expression in WCS365 with ColR-independent orthologs in PAO1; **Cluster 3**. Genes with ColR-dependent expression in PAO1 with ColR-independent orthologs in WCS365; **Cluster 4**. Genes with ColR-dependent in WCS365 without orthologs in PAO1; **Cluster 5**. ColR-dependent expression in PAO1 without orthologs in WCS365. Genes tested for rhizosphere colonization (Fig. 3) are marked with an asterisk.

Because only a small portion of differences in transcription in the rhizosphere were ColRdependent, we hypothesized that failure to induce expression of these genes in the rhizosphere may explain the requirement of ColR for rhizosphere colonization. We identified genes that were specifically upregulated in the rhizosphere in wild-type but not in a Δ*colR* mutant in both WCS365 and in PAO1 (Dataset S2; Methods). We identified 50 and 42 positively ColR-regulated genes with higher expression in the rhizosphere in wild-type than in the *colR* mutant in WCS365 and in PAO1, respectively (Fig. 2b, c). These data indicate that ColR may directly or indirectly upregulate a limited set of genes during rhizosphere colonization.

To determine if the ColR regulon is conserved across *Pseudomonas* spp., we performed comparative transcriptomics and genomics to identify orthologous genes present in the ColR regulons of PAO1 and WCS365^1^. We found that only 4 genes with ColR-dependent expression in the rhizosphere were orthologous between PAO1 and WCS365 (Fig. 2d). These include *eptA* and *eptC* (which encode phosphoethanolamine transferases), *tpbA* (which encodes a tyrosine phosphatase), and war*A* (which encodes a methyltransferase) (Fig. 2d; Cluster 1). Of these, *warA* was previously shown to be ColR-regulated in *P. putida*^30^. Despite ColR serving a similar function in rhizosphere colonization by *P. aeruginosa* PAO1 and *P. fluorescens* WCS365, these results suggest the ColR regulon may have largely diverged between PAO1 and WCS365.

Since the majority of genes in the ColR regulons of WCS365 and PAO1 did not overlap, we tested if the remaining ColR-dependent genes were part of the core *Pseudomonas* genome with strain-specific regulation, or if they were components of the *Pseudomonas* accessory genome. We surveyed the presence/absence of the remaining ColR-dependent genes across the genus *Pseudomonas*. We found that 26 ColR-dependent genes in *P. fluorescens* WCS365 and 21 ColRdependent genes in *P. aeruginosa* PAO1 had a ColR-independent ortholog in the other strain [Fig. 2d (Cluster 2 and 3) and Fig. S2], indicating the genes are conserved but have divergent regulation. The remaining genes in the ColR regulon encoded genes unique to PAO1 or WCS365; 20 ColR-dependent genes in WCS365 and 16 genes in PAO1 did not have any orthologs in the other strain (Fig. 2d; Cluster 4 and 5). This indicates that the divergence of the ColR regulon between strains may be due to both gain and loss of accessory genome components and through promoter evolution.

### Deleting components of the ColR regulon revealed strain-specific, but not host-specific, requirements for ColR-dependent genes

We reasoned that orthologous ColR-regulated genes shared in both *P. fluorescens* WCS365 and *P. aeruginosa* PAO1 might be part of a conserved ColR regulon required for host association. We tested if the *eptA*, *tpbA*, and *warA* orthologs, which are ColR-dependent in both *P. fluorescens* WCS365 and *P. aeruginosa* PAO1, were required for rhizosphere colonization in both strains. We used either transposon insertion mutants^31^ [PAO1 *tpbA::Tn5* and PAO1 *warA*::Tn5], or generated clean deletions (PAO1 Δ*eptA* and WCS365 Δ*eptA*, Δ*tpbA*, and Δ*warA*). We found that the *tpbA*::Tn5 mutant had reduced rhizosphere colonization in PAO1 but deletion of the WCS365 *tpbA* ortholog had similar rhizosphere colonization as wild-type WCS365 (Fig. 3a, b). While the WCS365 Δ*eptA* and Δ*warA* mutants had reduced rhizosphere colonization, the PAO1 Δ*eptA* and *warA*::Tn5 mutants did not (Fig. 3a, b). Collectively these data indicate that while ColR is a conserved determinant of rhizosphere colonization, ColR-dependent genes required for rhizosphere colonization have diverged between *P. fluorescens* WCS365 and *P. aeruginosa* PAO1.

**Figure 3.**
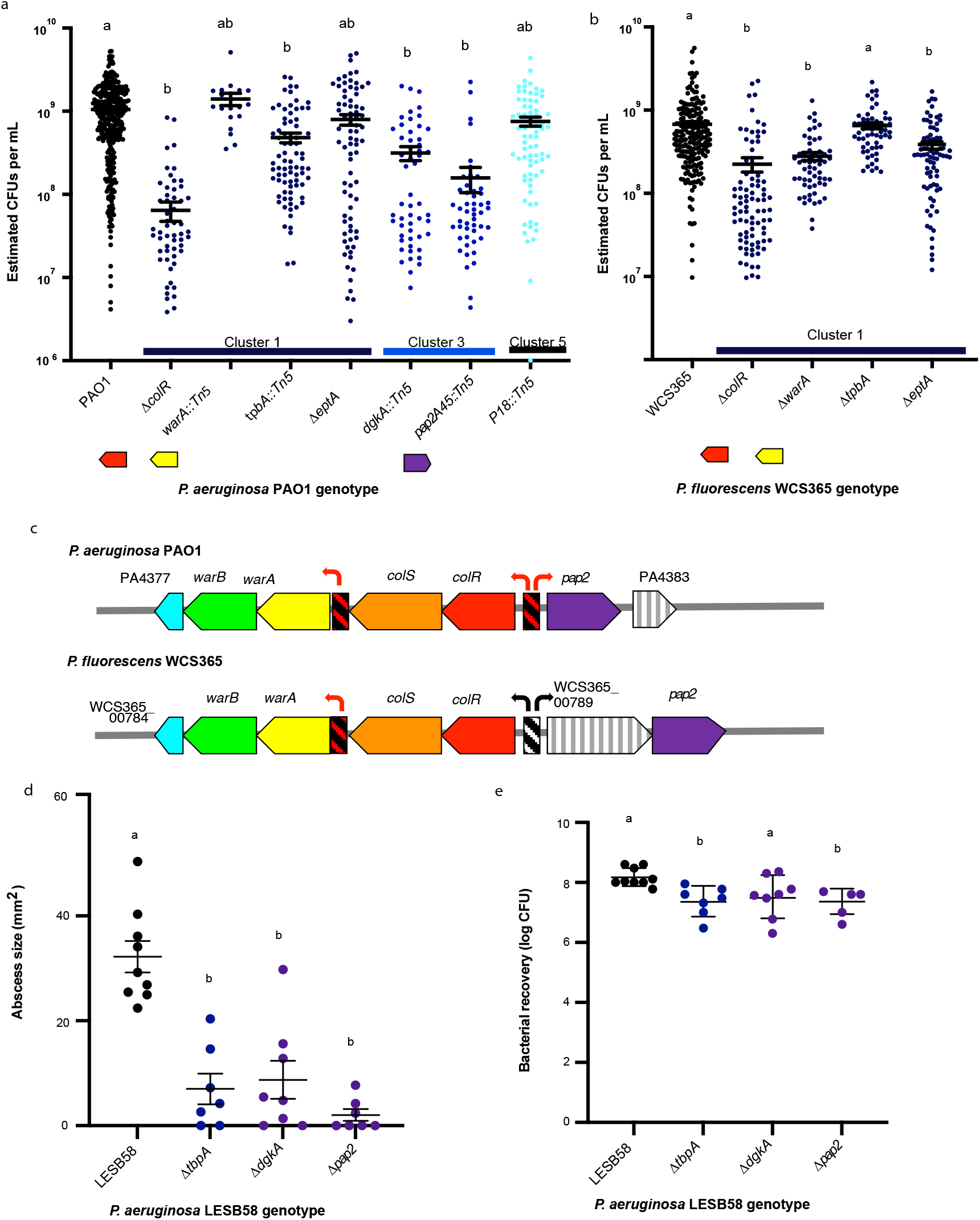
Strain-specific ColR-dependent genes are required for both rhizosphere colonization and virulence in a mouse abscess model. **(a)** Wildtype *P. aeruginosa* PAO1 or PAO1 mutants [Δ*colR*, *warA::Tn5* (PA4379), *tpbA::Tn5* (PA3885); Δ*eptA* (PA1927), *dgkA::Tn5* (PA3603), *pap2::Tn5* (PA4382), or PA4518::Tn5] and **(b)** Wildtype *P. fluorescens* WCS365 or WCS365 mutants [Δ*colR*, Δ*warA* (WCS365_00786), Δ*tpbA* (WCS365_00280), or Δ*eptA* (WCS365_03626)] containing GFP-expressing plasmids were inoculated into the Arabidopsis rhizosphere of hydroponically grown seedlings. Each data point represents the estimated cfu/mL for a single plant. Three independent experiments, each with a minimum of 10 plants, were performed (n > 30). Letters indicate significant differences at P < 0.05 using an ANOVA and Tukey’s HSD. Colours correspond to the gene clusters shown in Fig. 2d. **(c)** Genomic arrangement of homologous genes in *P. fluorescens* WCS365 and *P. aeruginosa* PAO1. Homologous genes are shown in the same color. Red arrows and striped, red boxes indicate putative ColR-binding sites and ColR-dependent transcription. Black arrows indicate ColRindependent transcription. **(d and e)** Wildtype *P. aeruginosa* LESB58 or LESB58 mutants [Δ*dgkA* (PALES_14321), Δ*tpbA* (PALES_10921), and Δ*pap2* (PALES_47610)] were injected into the subcutaneous thin skeletal muscle on the dorsum of mice. Abscess lesion size **(d)** and CFU counts **(e)** were determined 3 days post-infection. Error bars represent standard deviation. Letters indicate significant differences at P < 0.05 as determined by a Mann-Whitney U test.

To identify additional ColR-dependent genes that are unique to PAO1 and might be required for rhizosphere colonization or abscess formation, we made use of a transposon insertion library in *P. aeruginosa* PAO1^31^. We screened 24 PAO1 transposon insertion mutants, starting with insertions in genes with the greatest difference in expression between wildtype PAO1 and the Δ*colR* mutant in the rhizosphere (Dataset S2). To expedite screening, we measured OD_600_ to estimate bacterial growth in the rhizosphere 5 days after inoculation (Methods). We identified 10 PAO1 mutants that had a lower measured OD_600_ than wild-type PAO1 (a 3-fold decrease or lower), which were designated candidates for colonization deficiency (Dataset S2). We transformed 3 candidate mutants that exhibited the largest difference in expression between wild-type PAO1 and the Δ*colR* mutant (*dgkA*::Tn5,*pap2*::Tn5, and PA4518:Tn5) with a plasmid expressing GFP and measured fluorescence for more accurate quantification of bacterial growth in the rhizosphere (Fig. 3a). From this secondary screen we identified 2 additional mutants, *dgkA* (encoding a diacyl glycerol kinase) and *pap2* (with similarity to phosphatidic acid phosphatases), that had significantly decreased colonization in the rhizosphere (Fig. 3a).

Because we found that *colR* was required for virulence in mice (Fig. 1e), we investigated if *colR*-dependent genes required for *P. aeruginosa* PAO1 rhizosphere colonization were also required for virulence in the murine subcutaneous abscess model. We found that deletion of the *P. aeruginosa* LESB58 orthologs of the genes with rhizosphere colonization defects in PAO1 (*dgkA, pap2*, and *tpbA*), resulted in decreased virulence in the abscess model (Fig 3d). In addition, deletion of *tpbA* and *pap2* led to a small but significant decrease in bacterial recovery (Fig 3e), indicating that virulence of these mutants was impaired. Collectively these data indicate that an overlapping set of ColR-dependent genes are required for *P. aeruginosa* to associate with diverse hosts.

### *Pseudomonas* strains undergo functionally similar ColR-independent transcriptional reprogramming in the rhizosphere

Because we found that the largest differences in gene expression were due to differences in bacterial growth condition (minimal media vs. rhizosphere, Fig. 2a) rather than differences in genotype, we wanted to identify rhizosphere stressors encountered by *Pseudomonas* spp. that might reveal why ColR is necessary for rhizosphere colonization. We queried categories of genes that are induced in the rhizosphere in wild-type bacteria (Fig. 4a,b) us ing gene ontology (GO) analysis. We found that in both *P. fluorescens* WCS365 and in *P. aeruginosa* PAO1, the global changes in regulation were largely similar between wild-type bacteria and a *colR* mutant in the rhizosphere when compared to minimal media (Fig. 4a, b; Dataset S3). In both *Pseudomonas* species, we observed enrichment of genes involved in detoxification and survival in a harsh environment. These include metabolism of antibiotics and organic compounds, and genes involved in detoxifying the environment and responding to a new environment (Fig. 4; Dataset S3). The upregulation of genes involved in stress tolerance in both WCS365 and in PAO1 suggested that to grow in the rhizosphere, bacteria must protect themselves against harmful rhizosphere components.

**Figure 4.**
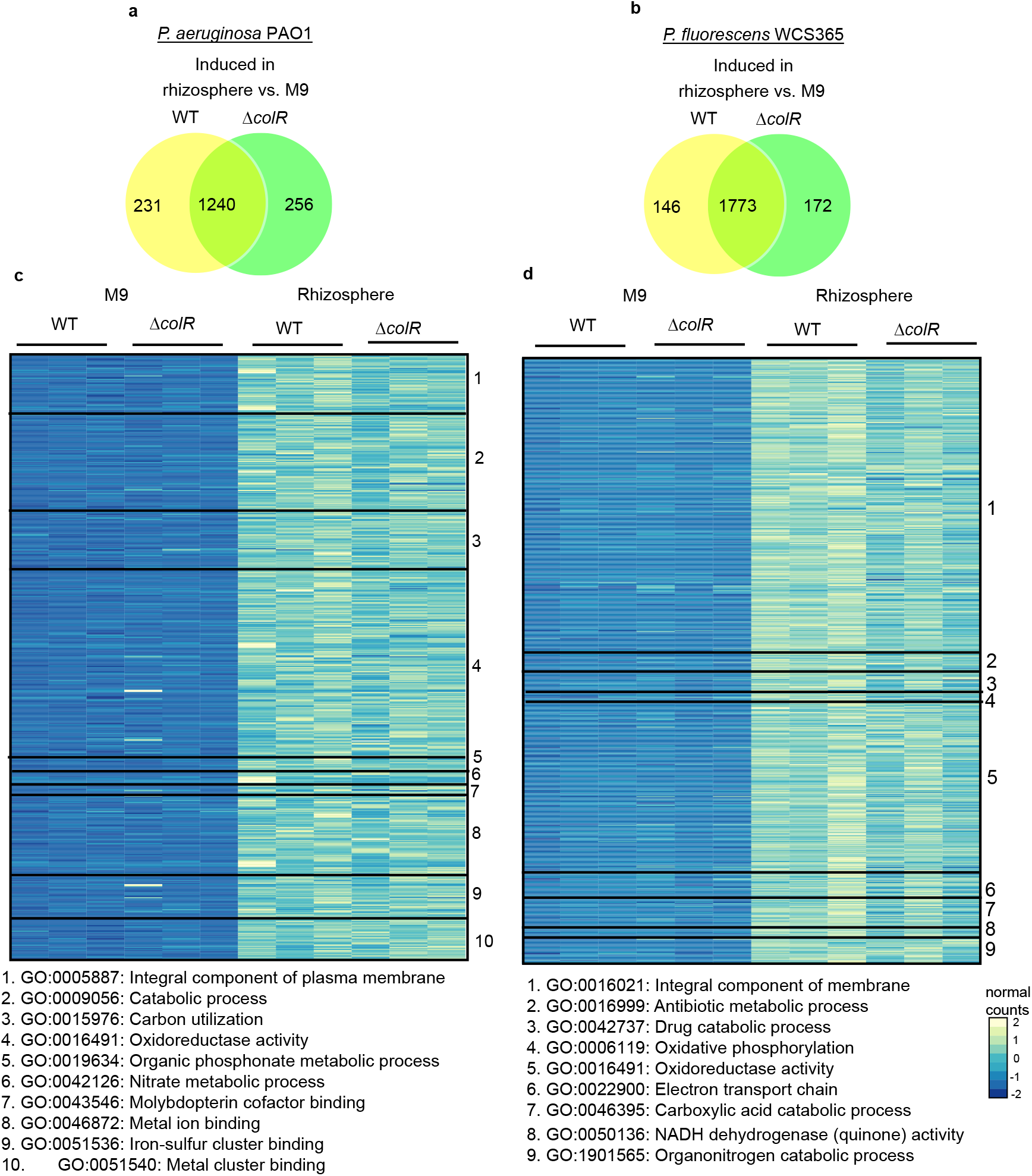
*Pseudomonas* undergo major ColR-independent transcriptional reprogramming in the rhizosphere. **(a and b)** Venn diagrams of the number of total genes in PAO1 **(a)** and WCS365 **(b)** that are significantly upregulated in WT or a *colR* mutant in the rhizosphere vs. M9 minimal media (greater than 0.585-fold change, padj < 0.01). **(c and d)** Significantly differentially expressed genes between the rhizosphere and M9 minimal media in wildtype PAO1 **(c)** and wildtype WCS365 **(d)**. Significantly enriched GO terms were identified using GOfuncR. Each number corresponds to a single parent GO term, as defined below the heatmaps.

Because we found upregulation of GO terms and processes related to protection against environmental stressors and membrane modifications, we tested if known rhizosphere stressors, including the presence of cationic peptides^32^, could explain the growth inhibition of a *colR* mutant. We found that the minimal inhibitory concentration (MIC) of the cationic peptide polymyxin B was similar between wild-type PAO1 and the *colR* mutant (Fig. S3). We also tested biofilm formation in a *colR* mutant, as biofilm formation was previously shown to be increased in a *colR* mutant and is important for virulence^33^ and rhizosphere colonization^34^. However, the *colR* mutant formed slightly lower levels of biofilm as wild-type bacteria in PAO1 and did not form lower levels of biofilm in LESB58 (Fig. S4). This could be due to the use of different *P. aeruginosa* isolates (PA14 vs. PAO1 and LESB58), or the different conditions under which biofilm formation was tested (endotracheal tube vs. 96well plate). These data indicate that the defect in rhizosphere growth of a *colR* mutant is not due to the presence of cationic peptides or a change in biofilm formation.

### ColR is necessary for growth at low pH and resistance to transition metals

We found that genes induced in the rhizosphere are enriched for GO terms including electron transfer activity and oxidoreductase activity (Fig. 4), suggesting that bacteria experience an ionic stress, such as low pH, in the rhizosphere. Previous studies have shown that the rhizosphere^35^ and microenvironments in the cystic fibrosis lung^36^ have low pH, leading us to wonder if low pH in a host environment inhibits growth of a *colR* mutant. However, we saw that growth of PAO1 Δ*colR* in LB at pH 5 was similar of that to wild-type PAO1 (Fig. 5b).

**Figure 5.**
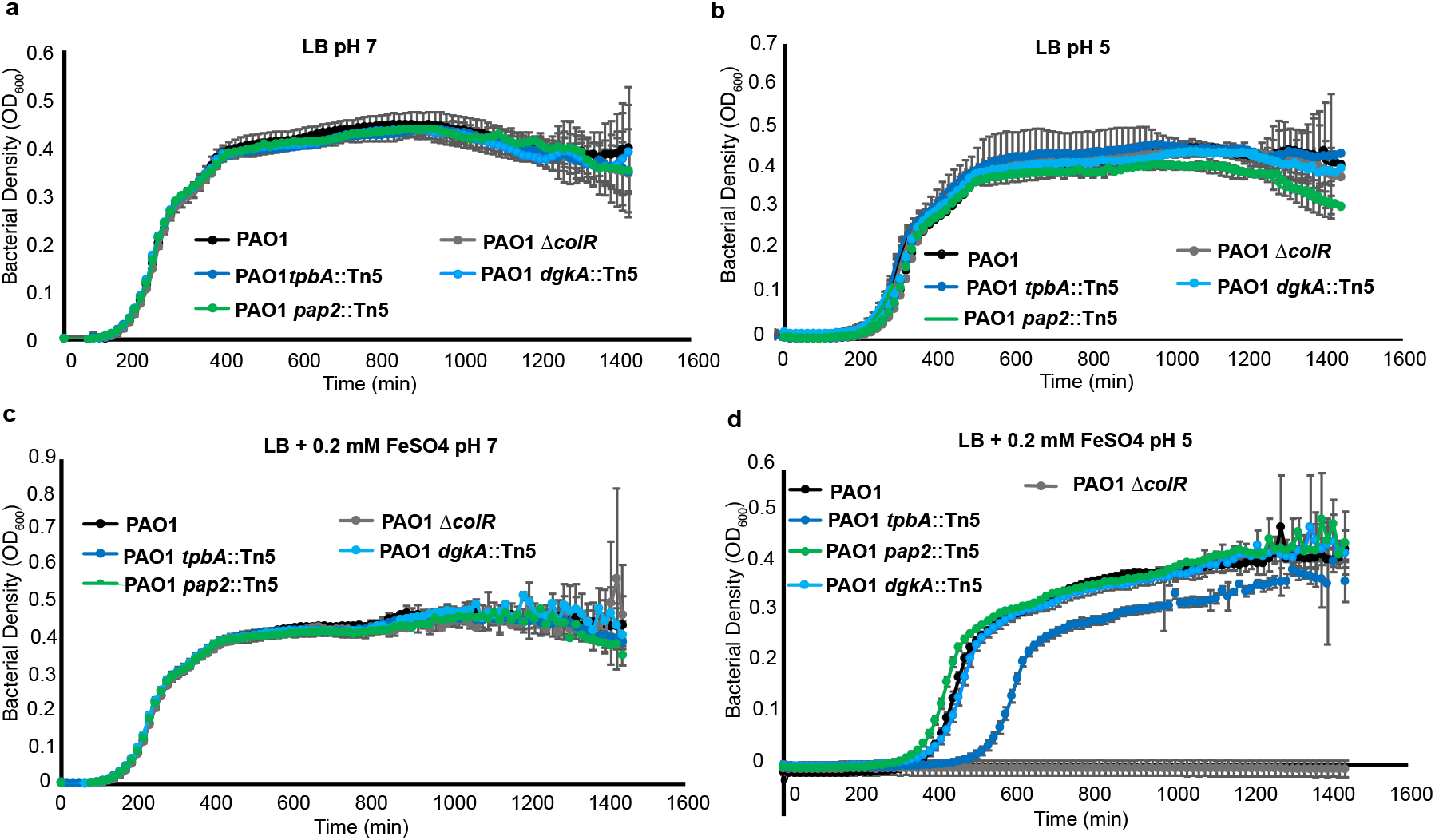
The *P. aeruginosa* PAO1 Δ*colR* mutant is sensitive to low pH and high iron, which is partially phenocopied by a *tpbA* mutant. Growth curves of wildtype PAO1 and PAO1 mutants (Δ*colR*, *tpbA::Tn5*, *dgkA::Tn5* or *pap2*::Tn5) in LB at pH 5 or 7 with and without the addition of 0.2 mM FeSO_4_. Growth curves were performed at least three times with similar results. Single experimental replicates are shown with averages and standard deviations of 6 technical replicates per time point.

Since genes encoding transition ion transporters for manganese (*mntP*), iron *fecA*), and copper (*copA1*) were dysregulated in the *colR* mutant compared to PAO1 (Dataset S2), we hypothesized that bacteria may experience metal stress in the rhizosphere. Accordingly, previous studies showed that a *P. putida colR* mutant is sensitive to zinc^30,37^. We found that growth of a *P. aeruginosa colR* mutant had increased sensitivity to high zinc concentrations, consistent with previous literature (Fig. S5). We determined the minimum inhibitory concentration (MIC) of four transition metals for PAO1 and the Δ*colR* mutant. We found that the *P. aeruginosa* PAO1 Δ*colR* mutant has lower tolerance to ZnSO_4_ (WT MIC = 10 mM; Δ*colR* MIC = 5 mM) and FeSO_4_ (WT MIC = 5 mM; Δ*colR* MIC = 2.5 mM) and similar tolerance to MnSO_4_ (WT MIC = 10 mM; Δ*colR* MIC = 10 mM) and CuSO_4_ (WT MIC = 10 mM; Δ*colR* MIC = 10 mM) than wild-type PAO1. Using metal concentrations below the MIC of *P. aeruginosa* PAO1 Δ*colR*, we performed growth curves with PAO1 and the Δ*colR* mutant and found that a PAO1 *colR* mutant has reduced growth relative to wild-type PAO1 in the presence of zinc, iron, and manganese (Fig. S6).

The concentrations of metals in MS media (30 μM Zn, 0.2 mM Fe, and 0.1 mM Mn), the media in which the plants are grown, is below the levels that inhibit the Δ*colR* mutant (3 mM Zn, 1 mM Fe, and 3 mM Mn); however, metal solubility increases at low pH so we wondered if a synergistic effect of pH and metal stress might be inhibitory^38^. We measured bacterial growth at pH 5 in LB the presence of 0.2 mM FeSO_4_ and found that *colR* is required for tolerance to this combination of metal and pH stress (Fig. 5d). We tested if *P. aeruginosa tpbA*::Tn5, *dgkA*::Tn5 and *pap2*::Tn5 mutants also had increased sensitivity to FeSO_4_ at low pH and found that *tpbA::Tn5* but not the other mutants partially phenocopied the growth defect of the Δ*colR* mutant (Fig. 5d). These data indicate that either that the loss of multiple ColR-dependent genes is required for sensitivity to the combination of 0.2 mM FeSO_4_ and low pH, or that loss of a ColR-dependent gene which we did not identify is required for this sensitivity. A mechanism of defence present in both plants and animals is the generation of reactive oxygen species (ROS) using iron^39,40^ suggesting a potential shared environment across hosts that might require ColR.

## Discussion

Bacteria in the genus *Pseudomonas* can form symbiotic associations with a strikingly diverse range of hosts with outcomes ranging from mutualism to pathogenesis. We showed that ColR is a conserved symbiosis factor, required for host-association not only in diverse *Pseudomonas* species, but also in diverse hosts. Interestingly, we found that the ColR regulon, and the ColR-dependent genes required for host association in distinct *Pseudomonas* spp., are largely divergent. Only 4 orthologous genes, *eptA, eptC, warA*, and *tpbA*, are ColR-dependent in both *P. fluorescens* and *P. aeruginosa*. We found that none of these genes are required for rhizosphere colonization in both species. This indicates that ColR/S may play a conserved role in sensing the host and responding through changes in transcription; however, the specific transcriptional changes depend on the bacterial genetic background.

We performed transcriptional profiling and GO analysis of genes upregulated in the rhizosphere in comparison to M9 minimal media, to identify genes that are likely involved in adaptation to growth in the rhizosphere environment (Fig. 4). This rhizosphere environment is composed of a variety of plant compounds, including specialized metabolites such as lignins, glucosinolates, antimicrobial peptides and indole compounds, and primary metabolites such as amino acids and sugars^41,42^. In both *Pseudomonas* species in our study, our data showed that processes related to metabolism were upregulated in the rhizosphere relative to minimal media, consistent with previous reports that the nutrient composition of the rhizosphere is highly complex. This allowed us to predict stressors that bacteria may encounter during rhizosphere colonization.

We further found that a Δ*colR* mutant is sensitive to a combination of low pH and high iron levels, suggesting a mechanism by which the rhizosphere or an abscess might inhibit growth or virulence, respectively, of a *colR* mutant. Metals are more soluble at low pH^38^, and a mechanism of defence common to both plants and animals is the production of reactive oxygen species (ROS) using iron^39,40^. We therefore hypothesize that increased solubility of metals could lead to an increased ROS production, inhibiting growth of a Δ*colR* mutant. As *P. aeruginosa pap2* and *dgkA* mutants were impaired in rhizosphere colonization and virulence, but were not sensitive to low pH and high metals, this indicates that there must be an additional stress or process that ColR is required to adapt to in both the plant rhizosphere and in the murine abscess. This could be an inhibitory antimicrobial peptide, lack of an essential nutrient, or a heightened immune response in the plant rhizosphere or mouse tissue. Revealing how Pap2 and DgkA protect *P. aeruginosa* in the rhizosphere and murine abscess can reveal additional stresses present in a host environment, and how the ColR regulon predisposes *Pseudomonas* for being broadly host associated.

We found that loss of *P. aeruginosa dgkA* and *pap2* resulted in reduced rhizosphere colonization and virulence in a murine abscess model (Fig. 3d). DgkA and Pap2 convert diacylglycerol (DAG) to phosphatidic acid (PA), and PA to DAG, respectively^43,44^. Both PA and DAG are key intermediates and precursors to the synthesis of many major membrane phospholipids^44^. Additionally, DAG is generated as a result of phosphoethanolamine (pEtN) addition to Lipid A^45^. Interestingly, we identified 3 pEtN transferase enzymes that were ColR-dependent in PAO1 [*eptA* (PA1972), *eptA2* (PA3310) and *eptC* (PA4517)] and 2 in WCS365 [*eptA* (WCS365_03626) and *eptC* (WCS365_05619)]. This suggests that *dgkA* may be required for turnover of the resulting DAG and that loss of *colR* may result in decreased membrane structural integrity through disruption of phospholipid synthesis and turnover.

This study has found that *colR* is required for rhizosphere colonization in *P. fluorescens* and in *P. aeruginosa* and abscess formation in *P. aeruginosa* but not through the regulation of conserved genes. This suggests that ColS may be a conserved host-sensing protein that activates ColR to adapt to a host environment; however, the ColR regulon may have diverged according to bacterial lifestyle. Divergence of two-component system regulons has been previously shown to occur in members of the family *Enterobacteriaceae*, in which the PhoP response regulator targets were largely divergent among species^46^. Collectively this work sheds light on the mechanisms by which bacteria can rapidly adapt to the host they are associated with, and the lifestyle of the bacterium.

## Materials and Methods

### Bacteria strains, media, and growth conditions

Bacteria strains and plasmids are listed in Dataset S1. All transposon mutants were obtained from the PAO1 two-allele library^31^. Routine bacterial culturing was performed using LB broth or agar at 37°C (for *P. aeruginosa* PAO1 and LESB58 as well as *E. coli*) or at 28°C (for *P. fluorescens* WCS365). When appropriate, LB was supplemented with sucrose, Kanamycin (Km), gentamicin (Gm), carbenicillin (Cb), and/or Irgasan.

### Strain construction

*E. coli* DH5α was used for construction and maintenance of plasmids, and *E. coli* SM10 λpir was used for biparental conjugation with *P. aeruginosa* and *P. fluorescens*. Deletion mutants were generated using the two-step allelic exchange method^47–50^. Deletion constructs for PAO1 Δ*colR* (PA4318), Δ*eptA* (PA1972), LESB58 Δ*colR* (PALES_47611)*, ΔtpbA* (PALES_14321)*, ΔdgkA* (PALES_47610)*, Δpap2* (PALES_10931), WCS365 Δ*colR* (WCS365_00788), Δ*eptA* (WCS365_03626), Δ*warA* (WCS365_00786), and Δ*tpbA* (WCS365_02280) were generated using primers listed in Dataset S1. As the flanking regions have 100% sequence identity, the same deletion construct for Δ*colR* was used to delete *colR* from both PAO1 and LESB58. Approximately 700-800 bp of sequence flanking the target genes were amplified using PCR. The flanking regions were joined using overlap extension PCR. The PCR products were digested using the appropriate restriction enzymes, as listed in Dataset S1, Tab 1, and ligated into the multiple cloning site of pEXG2 suicide vector containing *sacB*^51^. After confirming the insert by Sanger sequencing, the vector was transformed into *E. coli* SM10 λpir and transformants containing the plasmid were selected for on LB agar plates supplemented with 15 μg/ml Gm. The plasmid was then transferred to *Pseudomonas* via conjugation into PAO1, LESB58, or WCS365, and integration into the genome through homologous recombination. Merodiploids were selected using Gm (50 μg/ml for PAO1, and 500 μg/ml for LESB58) and irgasan (25 μg/ml) for *P. aeruginosa*, or Gm (15 μg/ml) and nalidixic acid (10 μg/ml) for *P. fluorescens*. Counter-selection on LB plates with 10% sucrose was performed to select for loss of the plasmid, which can result in either no deletion (if the second recombination event occurs in the region that the first recombination event occurred, leading to a loss of the plasmid), or a deletion (if the second recombination event occurs at the flanking gene on the opposite side of the genes as the first recombination event occurred). PCR and Sanger sequencing was used to screen for and validate deletion mutants.

The *colR* complementation plasmids were created using the pBBR1MCS-5 plasmid as a backbone for PAO1 and pBBR1MCS-2 for WCS365. PAO1 *colR*, and WCS365 *colR*, and approximately 200 bp upstream of each gene, respectively, were amplified using primers listed in Dataset S1. Amplicons and plasmids were digested with the appropriate enzymes and then ligated together, and the resulting insert was confirmed by Sanger sequencing. The *colR*complementation strains were generated by electroporating PEXG2-PAO1colR (P*colR*) into the PAO1 Δ*colR* mutant, and PEXG2-WCS365colR (*WcolR*) into the WCS365 Δ*colR* mutant. Gm (PAO1) or Km (WCS365) were used for selection and maintenance of the plasmid. Wild-type and Δ*colR* with empty vector (EV) were generated by electroporation with the pBBR1MCS-5 for PAO1 and pBBR1MCS-2 for WCS365.

GFP-expressing *P. aeruginosa* and *P. fluorescens* were generated by electroporating strains with pSMC21 (*Ptac-GFP*)^52,53^. Electrocompetent cells were prepared by pelleting, washing and resuspending cultures in 300 mM sucrose. Transformants were selected on and maintained using Cb (300 μg/ml) for *P. aeruginosa*, and Km (25 μg/ml) for *P. fluorescens*.

### Plant growth conditions

Axenic plants were generated by surface sterilizing Arabidopsis (wild-type accession Col0) seeds in 70% ethanol for approximately 2 min, 10% bleach for approximately 2 min, and washed three times with sterile water (H_2_O). Seeds were resuspended in H_2_O or 0.1% phytoagar and stored at 4°C in the dark for at least 48 h before sowing on Murashige and Skoog (MS) liquid media^54^, as described below. Plants were grown at 16 h light/8 h dark under 100 μM cool white fluorescent lights at 22°C.

### Rhizosphere colonization in 48-well plates

Plants were grown hydroponically in flat-bottom 48-well plates as described^17,27^. Briefly, one autoclave-sterilized Teflon mesh disk was placed in each well of a 48-well plate containing 250-300 μl MS plant growth media supplemented with 2% sucrose at pH 5.8. Sterilized Arabidopsis seeds were placed individually in the center of each disk. The plant media was changed to 270 μl of ½X MS media at pH 5.8 without sucrose after 10 days. At 12 days, overnight cultures of GFP-expressing bacteria were diluted to an OD_600_ of 0.0002 in 10 mM MgSO_4_, and 30 μl was inoculated into each well. GFP fluorescence (485 nm/535 nm excitation/emission) was read from the bottom of the wells each day up to 5 days post-inoculation, using a SpectraMax i3x fluorescent plate reader (Molecular Devices). Bacterial growth on a minimum of 10 plants was measured for each strain per experiment.

Bacterial abundance in the rhizosphere was estimated from the GFP fluorescence readings and reported in CFU per ml by first converting GFP fluorescence readings to OD_600_ using a standard curve, and then converted to CFU per ml using known values of number of bacteria per OD_600_ for both *P. fluorescens* WCS365 and *P. aeruginosa* PAO1. Standard curves of GFP fluorescence vs. OD_600_ were constructed for *P. aeruginosa* PAO1 and *P. fluorescens* WCS365. Separate standard curves were constructed for PAO1 and WCS365 strains that harbour two plasmids (pSMC21 and pBBR1MCS-5 or pBBR1MCS-2), because of possible differences in plasmid copy number due to the presence an additional expression vector.

### Study approval and animals

Animal experiments were performed in accordance with the Canadian Council on Animal Care (CCAC) guidelines following approval by the University of British Columbia Animal Care Committee (A14-0253). Mice used in this study were outbred CD-1 mice (female, 7-8 weeks). All animals were purchased from Charles River Laboratories, Inc. (Wilmington, MA). Mice weighed 25 ± 5 g at the experimental endpoint. Animals were group housed in cohorts of 4-5 littermates exposed to the same pathogen. Littermates were randomly assigned to experimental groups and standard animal husbandry protocols were employed.

### Murine subcutaneous abscess infection model

The role of ColR in virulence during high density infection was examined using *P. aeruginosa* LESB58)^55^ in a murine subcutaneous abscess model (Pletzer et al., 2017)^26^. Briefly, the LESB58 wild-type and mutant strains (Δ*colR*, Δ*dgkA*, Δ*pap2*, Δ*tpbA*) were subcultured at 37°C with shaking (250 rpm) to an OD_600_ = 1.0 in LB. Cells were washed twice with sterile phosphate buffered saline (PBS) and resuspended to a final OD_600_ = 1.0. Bacteria were injected (50 μl) subcutaneously into the right or left dorsum of mice for an inoculum density of ~5.0 x 10^7^ CFU. Abscesses were formed for 72 h with daily clinical grading. At the experimental endpoint, mice caliper, and abscesses were harvested in PBS. Then abscess tissues were homogenized using a Mini-Beadbeater (BioSpec Products, Bartlesville, OK) for bacterial enumeration on LB following serial dilution. Two or three independent experiments containing 24 biological replicates each were performed (*n* = 6-12).

### *Pseudomonas* sp. WCS365 genome assembly

Genomic DNA from *Pseudomonas* sp. WCS365 was sequenced on an ONT MinION sequencer and an Illumina MiSeq. The short reads were used to correct errors in the long reads using ropebwt2and^56^ FMLRC^57^. The corrected long reads were assembled into a single, circular contig using Flye^58^ with an expected genome size of 6 Mb. The long reads were mapped back to the assembly with minimap2^59^ and used to polish the genome using Racon^60^. The *Pseudomonas fluorescens* WCS365 genome will be available upon publication.

### Bacterial RNA extraction and RNA-Seq

RNA from rhizosphere-growing and minimal media-growing *P. aeruginosa* PAO1 and *P. fluorescens* WCS365 was extracted for RNA sequencing and analysis. M9 minimal media supplemented with 24 mM L-glutamine (L-gln) (*P. aeruginosa* PAO1), or supplemented with 30 mM sodium succinate (*P. fluorescens* WCS365) was used as the minimal media control. Plants were grown hydroponically as described in the 48-well plate rhizosphere colonization assay. Wells containing 12-day-old plants were inoculated with the wild-type and *colR* deletion mutants of *P. aeruginosa* PAO1 and *P. fluorescens* WCS365. To obtain sufficient RNA for sequencing, 6 wells containing plants or minimal media were inoculated with a final OD_600_ of 0.2 for PAO1 and 0.1 for WCS365. Both plant and minimal media plates were incubated in the plant growth conditions described above for 6 hours. After 6 hours, 40 μl of media from each of 6 plant wells or media from 3 minimal media wells were pooled and stabilized in RNAprotect^®^ Bacteria Reagent (QIAGEN) before performing RNA extraction using RNeasy Mini Kit (QIAGEN). When necessary, RNA was concentrated using ethanol precipitation.

RNA from three biological replicates was used with RNA integrity numbers (RIN) >9.9. cDNA library preparation and RNA-sequencing, using paired-end 150 bp reads, were performed by GENEWIZ using Illumina HiSeq. The sequencing yielded an average of 16,897,599 (standard deviation = 429,191) reads for each sample for all PAO1 and PAO1 Δ*colR* samples, and an average of 16,055,232 (standard deviation = 760,587) reads for each sample for all WCS365 and WCS365 Δ*colR* samples. The quality of the reads was assessed using FastQC v.0.11.8.^61^ Salmon v.1.1.0^29^ was used to align reads the PAO1 transcriptome and to obtain the count files for each sample. Reads were mapped to the PAO1 transcriptome with an average mapping rate of 70.8% (standard deviation = 1.3%) for each sample for PAO1 and were mapped to the WCS365 transcriptome with an average mapping rate of 75.5% (standard deviation = 3.5%) for each sample for WCS365. DESeq2 v1.26.0 was used for differential expression analysis in R^62^. Log_2_ fold change values were corrected using the *apeglm* package in R^63^. Genes with a log_2_ fold change of ≥ 1.5 (± 0.585 fold change) and adjusted p-value (padj) ≤ 0.01 were considered significantly differentially expressed. Normalized reads for each gene were generated using the DESeq2 package in R. The dplyr package was used for data manipulation (sorting and merging lists). The heatmap function in R was used to generate a heatmap using reads normalized for length of each gene. The *Pseudomonas fluorescens* WCS365 and *Pseudomonas aeruginosa* fastq RNAseq output files are available at NCBI GEO Accession no. GSE190448.

### Comparative Genomics

Orthology groups between the ColR-dependent genes in WCS365 and PAO1 were identified using the PyParanoid pipeline described previously^1,64^. Briefly, the PyParanoid pipeline was previously used to create a database using 3886 Pseudomonas genomes, identifying 24066 homologous protein families (“gene groups”) that covered 94.2% of the generated Pseudomonas pangenome. Groups shown in Fig. 2 were identified based on the PAO1 or WCS365 amino acid sequences and mapped onto a phylogenetic tree. Phylogenetic trees were generated using the PyParanoid pipeline as described previously^1,64^.

### Functional analysis of RNA-seq results

Gene ontology (GO) terms assigned to WCS365 were assigned using Blast2GO^65^. Lists of genes significantly differentially expressed in each condition or genotype were created using DEseq2, as described above. Lists of genes with assigned GO terms were inputted into R. GOfuncR (Gote, 2021) was used to identify GO terms significantly enriched in the conditions inputted (wild-type bacteria or Δ*colR* in the rhizosphere or minimal media). The go_enrich command was used to identify significantly enriched GO terms and GO terms with a FWER (family-wise error rate) ≤ 0.1 were refined to include only those terms with a refined p-value ≤ 0.05. Code is available on the Haney Lab Github page: https://github.com/haneylab/ColR_paper_RNAseq_analysis

### *In vitro* growth assays

For all growth curves, bacterial cultures were pipetted into clear, flat-bottom 96-well plates to a final OD_600_ of 0.02 and a final volume of 100 μl. Growth was monitored using a shaking plate reader (Molecular Devices VersaMax microplate reader), at a fixed temperature of 37°C for PAO1. OD_600_ readings were taken once every 15 min for 24 h. At least 4 technical replicates were used for each strain and condition. The growth curves for each combination of strain and condition were repeated 3 times.

Growth in FeSO_4_, ZnSO_4_, MnSO_4_, and CuSO_4_ were assessed by addition of the appropriate concentration of metals to LB. Metal concentrations used were below the MIC for both PAO1 and PAO1 Δ*colR*. To determine the MIC of the transition metals tested, LB media supplemented with 2X the highest concentration of metals (ZnSO_4_, MnSO_4_, FeSO_4_, and CuSO_4_) to be used for the MIC assay were prepared. The solutions of LB + metal were then serially diluted by 2X down the rows of round-bottom 96-well plates. The final volume in each well was 50 μl. Overnight bacterial cultures in LB were resuspended in fresh LB and diluted to an OD_600_ of 0.04. 50 μl of diluted bacterial cultures were added to each well. The 96-well plates were placed in an incubator at 37°C for 24 h before assessment of MIC. The MIC values were determined by eye and scored as the lowest concentration of metals in which there was at least a 50% reduction in growth.

Growth curves in ½X MS or M9 salts were performed as described above. L-gln at 24 mM was used as the carbon source in M9 minimal media. For growth curves at varying pH, ½X MS with 16 mM L-gln or 20 mM succinate was adjusted to pH 5, 6, or 7 using KOH. Overnight bacterial cultures in LB were washed and resuspended in 10 mM MgSO_4_ and diluted to an OD_600_ of 0.2. 90 μl of sterile media were aliquoted into each well in a 96-well plate before inoculation with 10 μl of the diluted bacterial suspension.

### Polymyxin B MIC assays

The MIC assay for PAO1 was performed according to a modified MIC method for antimicrobial peptides (http://cmdr.ubc.ca/bobh/method/modified-mic-method-for-cationic-antimicrobialpeptides/). Briefly, individual bacterial colonies grown on MH agar media were used to start overnight cultures in MH broth. Serial dilutions of polymyxin B were performed by making 20X solutions of the antibiotic in 0.01% acetic acid, 0.2% BSA and serially diluting the antibiotic twofold into sterile 96-well polypropylene microtitre plates. Cultures were then diluted to 5 x 10 ^6^ cfu/mL (OD_600_ = 0.0073) and 100 μL of this diluted culture was added to each well. The MIC values were determined by eye and scored as the lowest concentration of polymyxin B in which there was at least a 50% reduction in growth.

### Biofilm assay

*P. aeruginosa* PAO1 strains were tested in 24 h biofilm assays quantifying total biofilm biomass following the procedure of Haney et al., 2021^67^. Briefly, bacteria from overnight cultures were diluted to an OD_600_ of 0.05 in Synthetic Cystic Fibrosis Media without ammonium chloride^68,69^ (PAO1 strains) or LB (LESB58 strains) and 100 μl were inoculated into 96 well plates. After static growth for 24 h at 37°C, the planktonic bacteria were rinsed off with deionized water. Biofilm biomass was stained with 105 μl 0.1% crystal violet and destained with 110 μl 70% ethanol. Lastly, absorbance was measured at 595 nm. Absorbance of each sample was blanked by subtracting the average absorbance of the sterility controls, and then normalized to the average absorbance of the wild-type and expressed as %wild-type biofilm. Statistical significance was determined using an unpaired t-test if comparing two values, or a one-way ANOVA followed by a Tukey’s multiple comparisons test to correct for multiple comparisons if more than one value was being compared. p-values <0.05 were considered significant.

### Zinc plate inhibition assays

Growth in the presence of high levels of zinc was determined by spotting serial dilutions of the strains of interest on LB-agar plates containing high levels of zinc (1.2 mM for WCS365, 3 mM for PAO1). After 24h of incubation at 28 or 37 °C, plates were scanned using an Epson V850 Pro scanner, and growth inhibition was determined visibly by identifying if less than 50% of wildtype growth was seen after 24 h or 48 h of incubation for PAO1 or WCS365, respectively.

## Data availability

The WCS365 genome assembly and annotation are have been deposited in the National Centre for Biotechnology Information BioProject database and will be available upon publication of the manuscript. The RNA-seq raw sequencing has been deposited in the National Center for Biotechnology Information Gene Expression Omnibus database under accession GSE190448.

## Code availability

The code used for RNA-seq analysis is available from the Haney laboratory GitHub repository https://github.com/haneylab/ColR_paper_RNAseq_analysis.

## Acknowledgements

This work was supported by a CIHR Grant (PJT - 169051), NSERC Discovery Grant (NSERCRGPIN-2016-04121), and a Canada Research Chair salary award to CHH. REWH was funded by a CIHR foundation grant (FDN-154287) and received salary support from a Canada Research Chair and UBC Killam Professorship. YZ was supported by an NSERC CGS-M and CLW was supported by an NSERC CGS-D award. DT is supported by an NSF postdoctoral fellowship in Biology (IOS-2010946). MA is supported by a Vanier graduate scholarship. DP was supported by a fellowship from the Michael Smith Foundation for Health Research. We thank Drs. Fred Ausubel and Alina Gutu for conversations that lead to the conception of this project.

## Author contributions

Conceptualization, C.W., Y.Z, D.P., R.E.W.H., and C.H.H.; Methodology C.W. and J.Z.; Formal Analysis, C.W., Y.Z., M.A., M.D., A.W., D.P., and C.H.H; Investigation, C.W., Y.Z., M.A., D.T., M.D., and D.P.; Resources, R.E.W.H and C.H.H.; Data Curation, C.W., Y.Z., A.W., and C.H.H.; Writing – Original draft, C.W., J.Z., and C.H.H.; Writing – Reviewing and Editing, all; Visualization, C.W., J.Z., M.A., D.T., D.P., and C.H.H.; Supervision, C.H.H., D.P., and R.E.W.H.; Funding Acquisition, C.H.H., and R.E.W.H

