## Supplementary Figures for "The ColR/S two-component system is a conserved determinant of host association across *Pseudomonas* species"

### Supplemental Figures

**a**

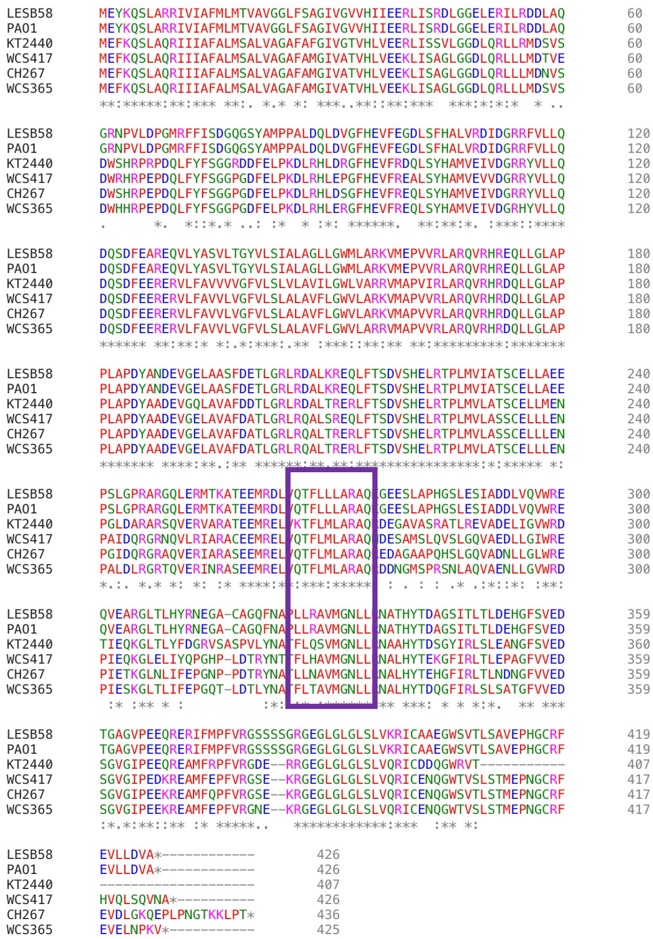

**b**

```
LESB58     MRILVVEDNRDILANLADYLSLKGYTVDCAQDGLSGLHLAATEHYDLIVLDVMLPGIDGY     60
PAO1       MRILVVEDNRDILANLADYLSLKGYTVDCAQDGLSGLHLAATEHYDLIVLDVMLPGIDGY     60
KT2440     MRILLVEDNRDILANLADYLGMKGYTVDCAQDGLSGLHLAATEHYDLIVLDIMLPGIDGY     60
CH267      MRILLVEDNRDILANLADYLGLKGYTVDCAQDGLSGLHLAATEHYDLIVLDIMLPGIDGY     60
WCS365     MRILLVEDNRDILANLADYLGLKGYTVDCAQDGLSGLHLAATEHYDLIVLDIMLPGIDGY     60
WCS417     MRILLVEDNRDILANLADYLGLKGYTVDCAQDGLSGLHLAATEHYDLIVLDIMLPGIDGY     60
           ****:***************.:*****************************:********

LESB58     ALCRRLREDARRDTPVIMLTARDQLDDRLQGFRSGADDYLVKPFALSELSARIEAVLRRA     120
PAO1       ALCRRLREDARRDTPVIMLTARDQLDDRLQGFRSGADDYLVKPFALSELSARIEAVLRRA     120
KT2440     TLCKRLREDARRDTPVIMLTARDQLDDRLQGFRSGADDYLLKPFALSELAARIEAVLRRA     120
CH267      TLCKRLREDARRDTPVIMLTARDQLDDRLQGFKSGADDYLVKPFALSELAARVEAVMRRT     120
WCS365     TLCKRLREDARLDTPVIMLTARDQLDDRLQGFKSGADDYLIKPFALSELAARIEAVMRRS     120
WCS417     TLCKRLREDARRDTPVIMLTARDQLDDRLQGFKSGADDYLLKPFALSELAARIEAVLRRA     120
           :**:******* ********************:*******:********:**:***:**:

LESB58     QGGGRRELSVADLSYDLDTLEVKRAGKSLKLNPIGLKLLAVLMQKSPHVVRRDALEEAVW     180
PAO1       QGGGRRELSVADLSYDLDTLEVKRAGKSLKLNPIGLKLLAVLMQKSPHVVRRDALEEAVW     180
KT2440     QGGGRRTLQVADLSYDLDTLEVTRQGRLLKLNPVGLKLLAVLMQKSPHVLRREVLEEALW     180
CH267      QGGGRRSLQVGDLSYDLDTLEVTREGKLLKLNPVGLKLLAVLMQKSPHVLRREILEEALW     180
WCS365     QGGGRRALQVGDLSYDLDTLEVTREGKLLKLNPVGLKLLAVLMQKSPHVLRREILEEALW     180
WCS417     QGGGRRALQVGDLSYDLDTLEVTREGRLLKLNPVGLKLLAVLMQKSPHVLRREILEEALW     180
           ****** *.*.***********.* *: *****:***************:**: ****:*

LESB58     GDDCPDSDSLRSHVHQLRQVIDKPFSVALLHTVHGVGYRLAEEPNGV*    227
PAO1       GDDCPDSDSLRSHVHQLRQVIDKPFSVALLHTVHGVGYRLAEEPNGV*    227
KT2440     GDDCPDSDSLRSHVHQLRQVIDKPFEKPLLHTVHGVGYRLAEGRDGV*    227
CH267      GDDCPDSDSLRSHVHQLRQVIDKPFAKPLLHTVHGVGYRLAEGRDGV*    227
WCS365     GDDCPDSDSLRSHVHQLRQVIDKPFDKPLLHTVHGVGYRLAEGRDGV*    227
WCS417     GDDCPDSDSLRSHVHQLRQVIDKPFAKPLLQTVHGVGYRLAEGRDGV*    227
           *************************   **:***********  :***
```

**Figure S1. ColR and ColS are highly conserved across diverse *Pseudomonas* species. (a)** Clustal Omega alignment of ColS amino acid sequences from 6 strains of *Pseudomonas* demonstrates that the protein is highly conserved. The predicted zinc binding site FExxE[37] boxed in purple is conserved among all 6 *Pseudomonas* strains. **(b)** Clustal Omega alignment of ColR protein sequences from the same 6 strains demonstrates high conservation of ColR across diverse *Pseudomonas* species. The DNA binding residues (boxed in purple) are 100% conserved across all strains.

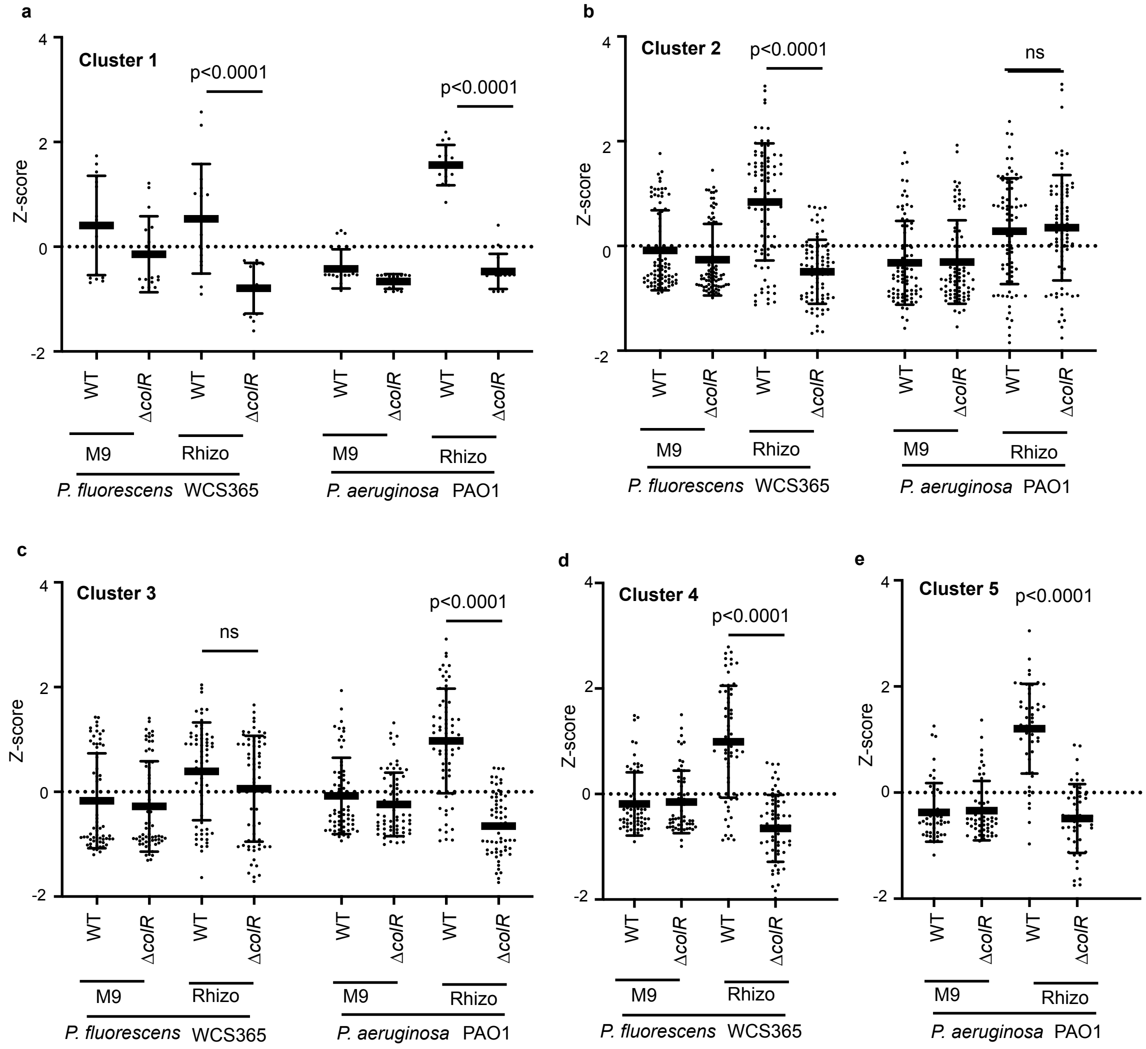


**Figure S2. Z-scores for ColR-dependent genes in each Cluster shown in Figure 2.** Genes fell into 5 distinct clusters: **(a) Cluster 1.** Genes with ColR-dependent expression in the rhizosphere in both WCS365 and PAO1; **(b) Cluster 2**. Genes with ColR-dependent expression in WCS365 in the rhizosphere with ColR-independent orthologs in PAO1; **(c) Cluster 3**. ColR-dependent expression in the rhizosphere in PAO1 with ColR-independent orthologs in WCS365; **(d) Cluster 4**. ColR-dependent in the rhizosphere in WCS365 without any orthologs in PAO1; **(e) Cluster 5**. ColR-dependent expression in the rhizosphere in PAO1 without any orthologs in WCS365. ANOVA followed by Tukey's HSD tests shows that orthologs of ColR-regulated genes in WCS365 (Cluster 2) and PAO1 (Cluster3) are not ColR dependent in the other strain.

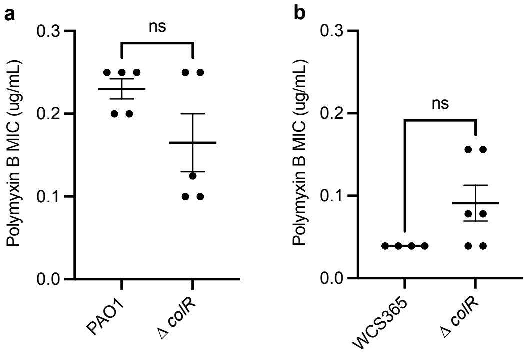


**Figure S3. Deletion of *colR* does not affect polymyxin B resistance in PAO1 or in WCS365.** Polymyxin B MIC assays reveal that the MIC of polymyxin B is not affected by the absence of *colR* in either **(a)** PAO1 or in **(b)** WCS365.

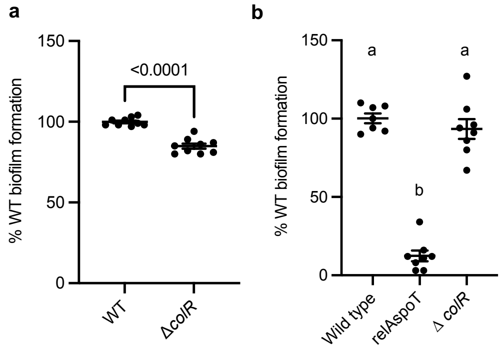


**Figure S4. Loss of *colR* does not consistently affect biofilm formation across different *P. aeruginosa* species.** (**a**) Loss of *colR* leads to a small, but significant, decrease in biofilm formation in PAO1. (**b**) In LESB58, a double deletion of genes previously identified to be requited for biofilm formation, *relA* and *spoT*[70], leads to a significant decrease in biofilm formation, but deletion of *colR* does not. In (**a**) p-value was determined by performing an unpaired t-test; in (**d**) significance was determined using a one-way ANOVA followed by a Tukey's HSD. P-values <0.05 were considered significant.

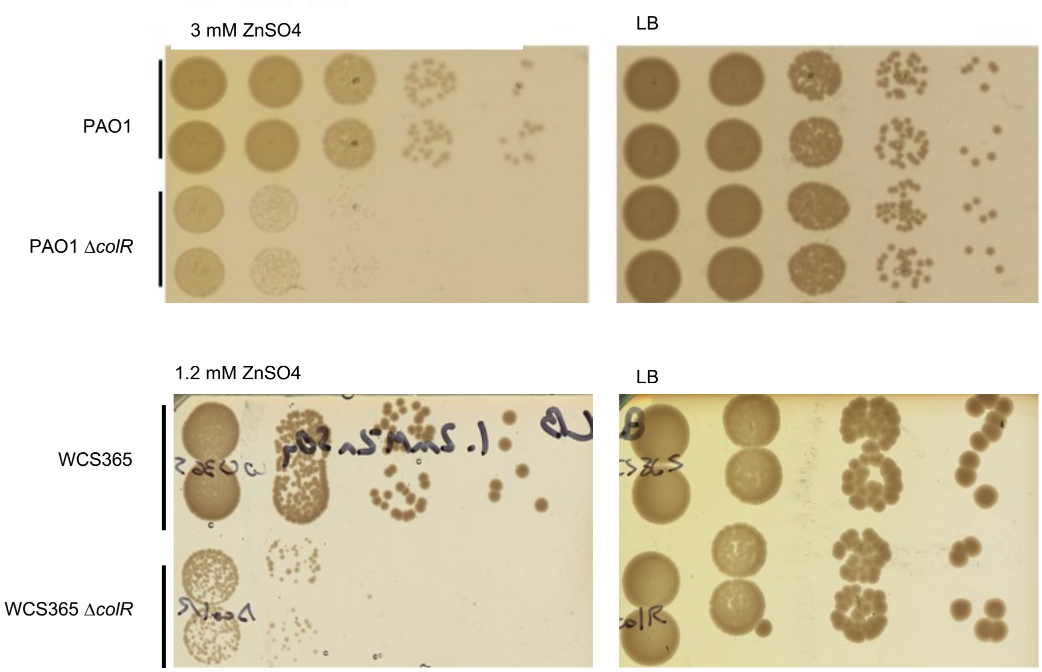


**Figure S5. Loss of *colR* leads to increased zinc sensitivity in both *P. aeruginosa* PAO1 and in *P. fluorescens* WCS365.** Deletion of *colR* in PAO1 leads to a visible decrease in growth on LB media containing 3 mM $ZnSO_4$ (top). Deletion of *colR* in WCS365 leads to a visible decrease in growth on LB media containing 1.2 mM $ZnSO_4$ (bottom).

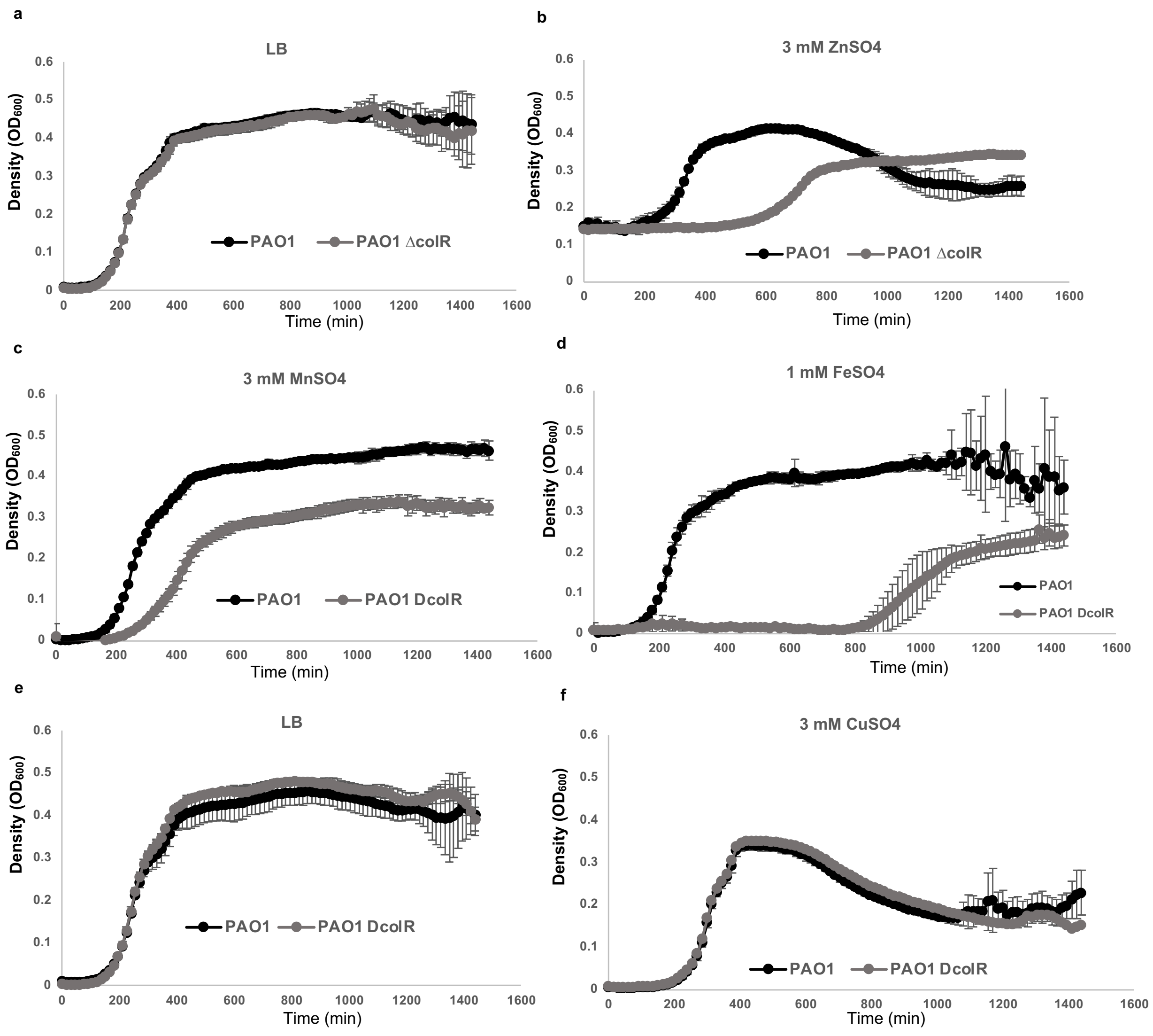


**Figure S6. Deletion of *colR* leads to increased sensitivity to zinc, manganese, and iron, but not copper, in PAO1.** Deletion of PAO1 *colR* leads to a longer lag phase and shorter doubling time than wildtype **(a and e)** in the presence of 3 mM $ZnSO_4$ (**b**), a longer lag phase, shorter doubling time, and lower stationary phase density in the presence of 3 mM $MnSO_4$ **(c),** and 1 mM $FeSO_4$ **(d),** but does not affect growth in 3 mM $CuSO_4$ **(f).** Graph **(a)** shows wildtype and *colR* mutant growth of bacteria in LB grown in the same 96-well plate as bacteria grown in zinc, magnesium and iron. Graph **(e)** shows wildtype growth and *colR* mutant growth in LB grown simultaneously with bacteria grown in copper. Error bars represent standard deviation. Graphs are representative of 3 independent growth curves each with 3-6 technical replicates per condition.
